# Maintaining the provenance of microscopy metadata using OMERO.forms software

**DOI:** 10.1101/109199

**Authors:** Douglas P.W. Russell, Peter K. Sorger

## Abstract

The creation of datasets that are findable, accessible, interoperable and reproducible (the FAIR standard) requires that data provenance be maintained^1^. Provenance is particularly important for microscopy data, whose interpretation is dependent on the biological context (e.g. cell state) and detection reagent (e.g. antibody.) This paper describes a new software tool, OMERO.forms, that extends the OMERO microscopy data management system^2^ to simplify and enhance metadata entry and provenance tracking.

---

Data provenance comprises a record of data origins, the context in which it was collected, and how both data and context evolved over time. It is particularly important to track the provenance of metadata in a collaborative system in which multiple individuals are involved in data collection, curation and analysis. Increasing adoption of management software for microscopy data and metadata presents an opportunity to develop new software for tracking data provenance. OMERO^2^ is the preeminent software system for managing microscopy data and metadata. OMERO is comprised of a repository server, an Application Programming Interface (API) and two standard clients: the OMERO.insight client is desktop based and OMERO.web is browser-based. The repository stores and organises image data and metadata on a remote server. The API enables interaction with all features of the repository, including: importing data, adding/querying/searching metadata and retrieving specific images and image planes (in the case of multi-dimensional images). However, even with within the context of a formal informatics system such as OMERO, managing metadata from microscopy experiments is challenging.^**3**^ Across an experiment, some metadata is relatively homogenous (e.g. instrument properties) but biological conditions or antibody identity often vary from one experiment to the next. This heterogeneous metadata must also be recorded, verified to be valid and ideally have its provenance tracked.

It is infeasible for OMERO to provide a pre-specified schema for every type of experimental metadata. Instead, OMERO has general purpose mechanisms for attaching metadata to an *imaging collection object,*^*4*^ a specific collection of data arising from an experiment, multi-experiment project or high content screen. One of the most useful metadata association mechanisms in OMERO is *map annotation*. *Map annotations* are key-value pairs: an identifier for the item of data (the key) and the actual value of the data (the value). Map annotations are flexible and can be used to store a wide range of metadata. OMERO APIs and clients can programmatically add and read annotations for a given object allowing annotations to be used to mark-up figures, convey key information to other investigators, and utilised by data analysis pipelines.

Although *map annotations* are potentially powerful, allowing unrestricted annotation of objects with them runs the very considerable risk of mistyped terms and omissions caused by human error. Information in *map annotations* is not checked for consistency against a schema, a formal model of metadata structure. For example, there is no mechanism for mandating the presence of a particular key, such as cell type, or ensuring that a value that is entered conforms to a particular type (number, text string etc.) or a constrained vocabulary (hence a proliferation in controlled metadata of nominally identical names such as SK-BR-3, SKBR3 etc.). OMERO.forms software addresses these issues directly.

OMERO.forms is an extension of OMERO.web that enhances metadata input and records data provenance. OMERO.forms exploits the fact that, while is it unrealistic for experimental metadata to be homogenised so that it fits a fixed schema across an entire research organization or field of science, it is usually the case that a specific set of experiments have similar metadata requirements. In OMERO.forms, experiment-specific information is entered through reusable, but customisable HTML forms that make sure that information entered conforms to the specified form design. A form provides users with a context for a particular class of metadata input and is used to render requirements on that metadata in an accessible way, such as presenting standardized vocabularies as a dropdown selection. Both information submitted by a user and the form design itself (including its formal schema) are stored within OMERO in an immutable metadata store to ensure the integrity of data provenance. There is no limit to the number of new or updated form submissions that can be made. Thus, OMERO.forms combines the advantages of flexible map annotations with the constraints of formalized data schemas.

New forms are created using a web-based client in OMERO.forms and are associated with one or more types (e.g. dataset or high content screen) of *imaging collection object.* As forms are being created, the design is rendered live in HTML and can be tested by entering and validating exemplar metadata. Forms designed and presented with OMERO.forms have the capability to restrict a field to a certain type of metadata (string, number, boolean, etc.) or a controlled vocabulary (e.g. Male/Female). These fields can be a single value or a list of values of the specified type or vocabulary. Fields can optionally be made mandatory. Forms can also be designed with help text to guide user input and provides feedback on input that does not validate against the form design. The feedback is visualized in the context of the field or fields in which the validation failed. Form designs are stored as JSON (JavaScript Object Notation) conforming to JSON Schema.^5^ The JSON Schema conformance is what allows OMERO.forms to validate information entered into the form against the form’s schema (i.e. the form design) and to render the JSON designed form into an HTML form for display. A form is used throughout the course of a particular study to manage metadata entry and maintain provenance; as many different form schemas as are needed can be created for the types of experiments being conducted. The web application has an interface to allow OMERO group owners to assign forms of interest to the specific groups for which they are relevant. This is useful in systems with many forms as it ensures that users are only presented with forms that are applicable to them.

A key feature of OMERO.forms is that it presents metadata in an intelligible form and also records the entire provenance of that metadata, including initial user input and any revisions. When viewing, or editing a form for which data has previously been entered, the most recent data is preloaded into the form for reference. Every form version ever submitted is retained by the system and information submitted through a form is associated with the specific version of the form that was used to enter it. This is important because metadata is often uninterpretable when divorced from the context in which it was collected. An interface for viewing the full provenance of a form is available (Fig. 1a): all previous metadata submissions for a specific form are presented as a list along with a time of creation timestamp, username of the editor and any change messages (descriptions of the changes made) submitted. Clicking on any of these entries renders the metadata in the context of the form in which it was originally entered. Moreover, when new information is submitted to a field in in OMERO.forms, the metadata is stored as a JSON encoded string to a metadata store that is not accessible (and thus not changeable) to regular OMERO users. The submission is stored in the context of the imaging collection object and form for which it was submitted. Previous metadata submissions are not overwritten and the chronology of these represents the history of that form metadata. In addition to the immutable store, metadata is encoded as map annotations (Fig. 1b) on the relevant imaging collection object. No special knowledge of OMERO.forms is required for other software to make use of this annotation since annotating an imaging collection object is a standard functionality of the core OMERO system itself. Thus, all metadata is searchable and easily visualised in existing OMERO clients and available for use in computational analyses through existing OMERO APIs.

**Figure 1:**
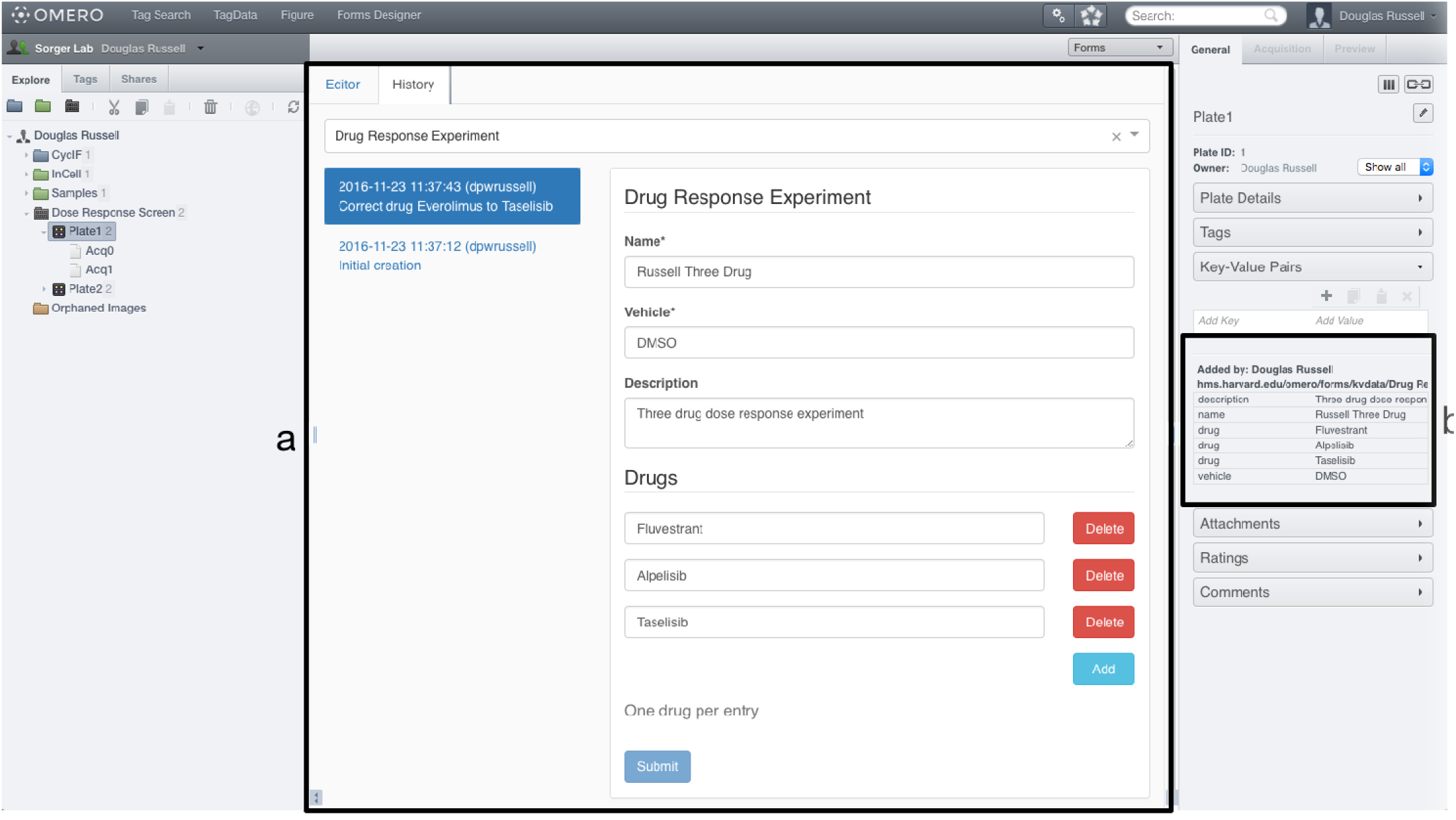
OMERO.forms History view. {a} History, {b} Associated map annotation. **Fig. 1a** The history viewer in OMERO.forms. Shows the information from every submission ever made for the given form and imaging collection object rendered in the context of the form version for which it was submitted. **Fig. 1b** The default OMERO visualization of a map annotation showing key-value pairs. This is attached directly to the imaging collection object on form submission to allow OMERO and any OMERO compatible software to access it without explicit knowledge of OMERO.forms.

OMERO.forms addresses key requirements for application of FAIR standards to image data while requiring little or no modification of existing clients and APIs. OMERO.forms is free, open source software, available on GitHub^6^ that can be installed on any OMERO server v5.0 or greater. The client and interface are intuitive enough for non-expert users to create new forms, enter metadata and review provenance while also including management features for group owners.

## Acknowledgements.

This work was supported by the NIH LINCS and BD2K programs; grants U54 HL127365 and 3U54HL127624-03S2, and HMS Research Computing.

## Methods

Immutable metadata store. OMERO has no built-in immutable data store that a user can make use of. OMERO.forms makes use of privilege escalation in OMERO.web to record form metadata and context as an OMERO map annotation attached to a special user created for OMERO.forms. The annotations attached to this “formmaster” user are not readable or writeable by regular users. To access a forms history, privilege escalation is again used. At the point of privilege escalation, authorization checks are performed to ensure that a user has the right to access the requested form data.

JSON Schema. An IETF (Internet Engineering Task Force) draft specification^5^ for validating schemas written in JSON. An implementation of this specification underpins OMERO.forms. This implementation is used to render an HTML form from the JSON design of a form. It is also used to validate values entered into the rendered form as conforming to assertions set forth in the design.

OMERO.forms Designer. This part of the web application is used to design forms. Supplementary Fig. 1 shows the form editor interface. The section in supplementary Fig. 1a is used to give a form a unique name and configure which OMERO object types it is applicable to. Previously created forms can be loaded in this section and if the editing user has the permission to do so, the loaded form can be edited and saved. If the user does not have permission to (or does not want to) update an existing form, they can always save it (with or without modifications) with a new name. This section is also used to enter the change message when creating or updating a form which can be useful in retrospect to understand why changes were made. Supplementary Fig. 1b, supplementary Fig. 1c and supplementary Fig. 1c are html *textarea* input boxes enhanced with some basic syntax highlighting, automated indenting and line numbers. Supplementary Fig. 1b and supplementary Fig. 1c are used to do the actual form design. Supplementary Fig. 1b designs the schema of the form including the names, order, defaults and restrictions of the elements and their types. A simple named string field “vehicle” is shown. Supplementary Fig. 1c customises the visual representation of the form. In this example the “description” field (simply a string) would default to a text input (a single line of text), but has been customised to show a textarea (multiple lines of text) by changing the “widget” used to display it. A large number of widgets are built-in and this is extensible to allow the addition of custom higher-order widgets as the need arises. Supplementary Fig. 1d allows the manual input of test data and also shows a live rendering of data entered into the currently displayed form preview. Supplementary Fig. 1e is a live and functional rendering of the form and will update as changes are made to the form’s design. This is exactly how the form will appear to the user at the point of use including the display of errors for invalid/omitted fields.

**Supplementary Figure 1:**
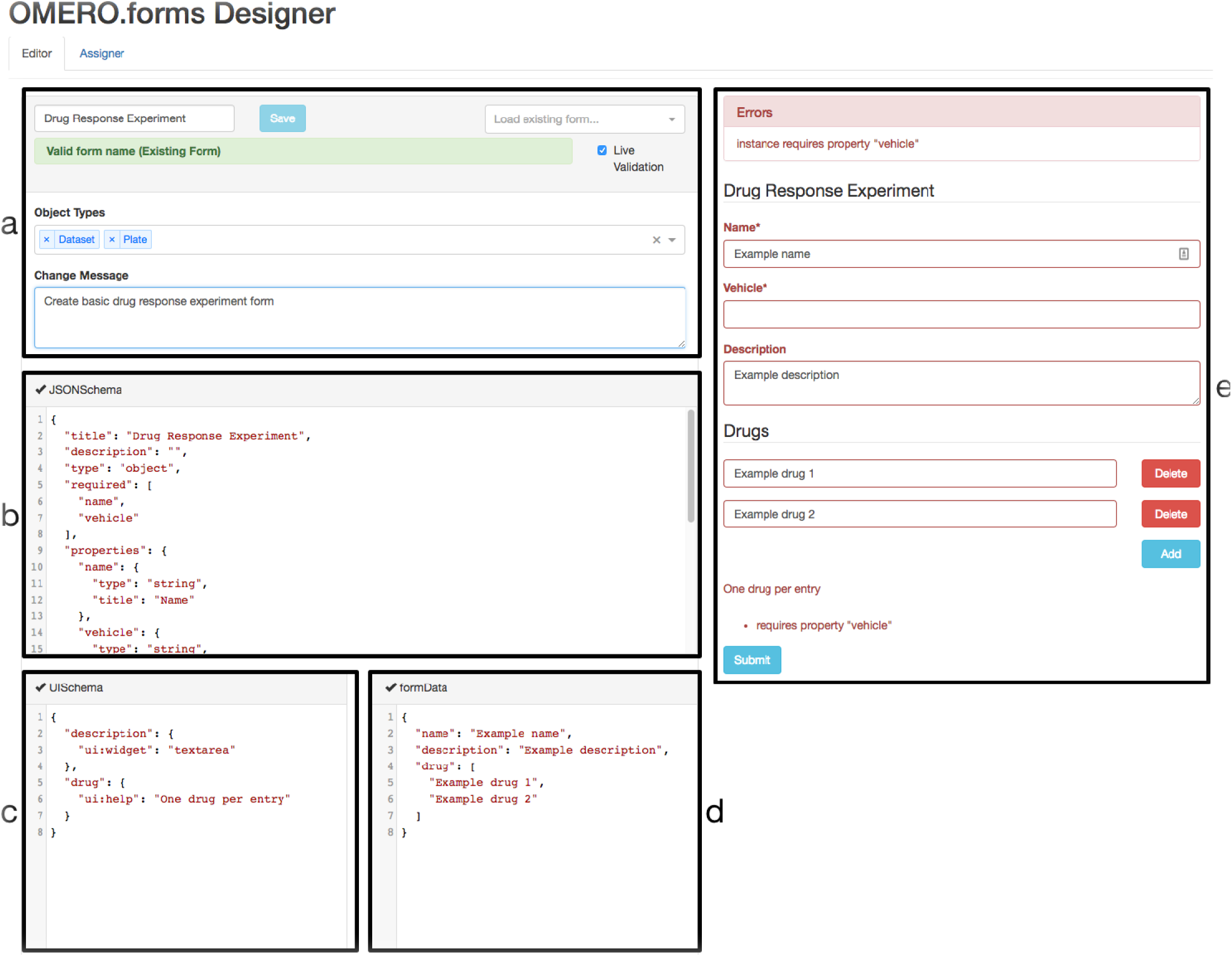
OMERO.forms Designer. {a} Form configuration, {b} Form design, {c} Form UI (User Interface) design, {d} Form data, {e} Form preview **Supplementary Fig. 1a** OMERO.forms designer configuration of a form detailing the imaging collection object types it is applicable to, a name and an opportunity to enter a message describing what changes have been made to the form in the current update. **Supplementary Fig. 1b** The JSON representation of the form schema. The form design is changed by modifying this. **Supplementary Fig. 1c** The JSON representation of the form UI schema. Customization of how the HTML form is rendered from the JSON form schema is configured here. **Supplementary Fig. 1d** The data structure representing the information entered into the rendered form. This can also be used to enter test data directly to see how it is rendered in the context of the current rendered form. **Supplementary Fig. 1e** The live rendering of the form schema as an HTML form.

Usage of OMERO.forms for end-users. Making use of OMERO.forms as an end-user is very simple as demonstrated in supplementary Fig. 2. When selecting an imaging collection object in OMERO.web (Supplementary Fig. 2a), choose the “forms” centre panel option from the dropdown box (Supplementary Fig. 2b). This will display the OMERO.forms interface with a list of all forms available to the current user. Upon selecting a form (Supplementary Fig. 2c), the form is rendered in the centre panel. If this form has been previously submitted for the given object, the most recent values will be loaded into the fields, otherwise they will be blank (or defaults if configured). The user can then populate the form and once the whole form is valid, submit it. The submitted form values are saved to the immutable data store and converted into a map annotation which is attached to the object and displayed by OMERO.web by default (Supplementary Fig. 2d).

---

**Supplementary Figure 2:**
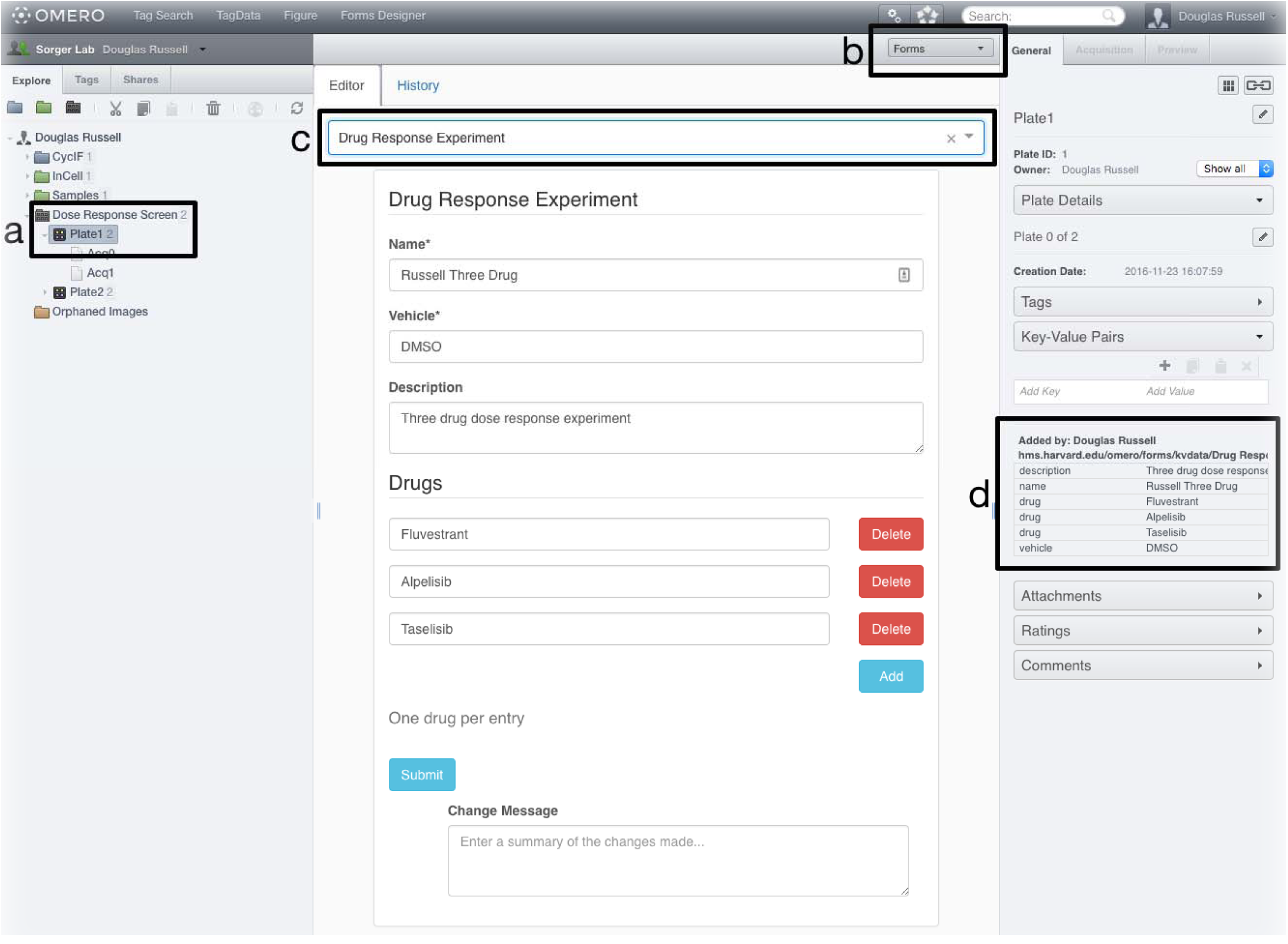
OMERO.forms Editor. {a} Object selection, {b} Plugin selection, {c} Form selection, {d} Associated map annotation **Supplementary Fig. 2a** The imaging collection object currently selected. In this case, a plate from a high content screen. **Supplementary Fig. 2b** The centre panel plugin currently selected. In this case, “Forms” to represent OMERO.forms. **Supplementary Fig 2c** The currently selection form. In this case a basic “Drug Response Experiment”. **Supplementary Fig 2d** The default OMERO visualization of a map annotation showing the data from the current form.

## References

1. M. D. Wilkinson, M. Dumontier, I. J. Aalbersberg, G. Appleton, M. Axton, A. Baak, N. Blomberg, J. W. Boiten, L. B. da Silva Santos, P. E. Bourne, J. Bouwman, A. J. Brookes, T. Clark, M. Crosas, I. D illo, O. Dumon, S. Edmunds, C. T. Evelo, R. Finkers, A. Gonzalez-Beltran, A. J. Gray, P. Groth, C. Goble, J. S. Grethe, J. Heringa, P. A. ’t Hoen, R. Hooft, T. Kuhn, R. Kok, J. Kok, S. J. Lusher, M. E. Martone, A. Mons, A. L. Packer, B. Persson, P. Rocca-Serra, M. Roos, R. van Schaik, S. A. Sansone, E. Schultes, T. Sengstag, T. Slater, G. Strawn, M. A. Swertz, M. Thompson, J. van der Lei, E. van Mulligen, J. Velterop, A. Waag-meester, P. Wittenburg, K. Wolstencroft, J. Zhao, and B. Mons. The FAIR Guiding Principles for scientific data management and stewardship. Sci Data, 3:160018, Mar 2016.

2. C. Allan, J. M. Burel, J. Moore, C. Blackburn, M. Linkert, S. Loynton, D. Macdonald, W. J. Moore, C. Neves, A. Patterson, M. Porter, A. Tarkowska, B. Loranger, J. Avondo, I. Lagerstedt, L. Lianas, S. Leo, K. Hands, R. T. Hay, A. Patwardhan, C. Best, G. J. Kleywegt, G. Zanetti, and J. R. Swedlow. OMERO: flexible, model-driven data management for experimental biology. Nat. Methods, 9(3):245–253, Feb 2012.

3. J. R. Swedlow, I. G. Goldberg, and K. W. Eliceiri. Bioimage informatics for experimental biology. Annu Rev Biophys, 38:327–346, 2009.

4. S. Li, S. Besson, C. Blackburn, M. Carroll, R. K. Ferguson, H. Flynn, K. Gillen, R. Leigh, D. Lindner, M. Linkert, W. J. Moore, B. Ramalingam, E. Rozbicki, G. Rustici, A. Tarkowska, P. Walczysko, E. Williams, C. Allan, J. M. Burel, J. Moore, and J. R. Swedlow. Metadata management for high content screening in OMERO. Methods, 96:27–32, Mar 2016.

5. Austin Wright and Geraint, Luff. JSON Schema Validation: A Vocabulary for Structural Validation of JSON. Internet-Draft draft-wright-json-schema-validation-00, Internet Engineering Task Force, October 2016. Work in Progress.

6. D.P.W Russell. OMERO.forms. https://github.com/sorgerlab/OMERO.forms, 2016.

